# High-density spinal cord stimulation selectively activates lower urinary tract afferents

**DOI:** 10.1101/2021.04.30.442206

**Authors:** Maria K Jantz, Chaitanya Gopinath, Ritesh Kumar, Celine Chin, Liane Wong, John I Ogren, Lee E Fisher, Bryan L McLaughlin, Robert A Gaunt

## Abstract

Epidural spinal cord stimulation (SCS) has recently been reported as a potential intervention to improve limb and autonomic functions, with lumbar stimulation improving locomotion and thoracic stimulation regulating blood pressure. We asked whether sacral SCS could be used to target the lower urinary tract. Here we show that high-density epidural SCS over the sacral spinal cord and cauda equina of anesthetized cats evokes responses in nerves innervating the bladder and urethra and that these nerves can be activated selectively. Sacral epidural SCS always recruited the pelvic and pudendal nerves and selectively recruited these nerves in all but one animal. Individual branches of the pudendal nerve were always recruited as well. Electrodes that selectively recruited specific peripheral nerves were spatially clustered on the arrays, suggesting anatomically organized sensory pathways. This selective recruitment demonstrates a mechanism to directly modulate bladder and urethral function through known reflex pathways, which could be used to restore bladder and urethral function after injury or disease.

## Introduction

Lower urinary tract (LUT) dysfunction occurs in 20-40% of the global population^1^ and has an economic impact measured in billions of dollars in medical costs every year^2^. One common clinical problem is overactive bladder; people experience excessive bladder contractions that increase the frequency with which they feel the urge to void^3^. Overactive bladder reduces sleep quality and participation in daily activities, and is associated with increased incidence of urinary tract infections^3^. Furthermore, losing voluntary bladder control is one of the least visible but most limiting consequences of spinal cord injury (SCI), making improvements in bladder control one of the highest priorities for people with SCI^4^. Unfortunately, current treatment methods for people living with neurogenic bladder dysfunction, particularly catheters, only address symptoms and routinely cause urinary tract infections requiring hospitalization^5,6^.

Electrical stimulation of the nervous system offers the potential to address the underlying causes of neurogenic bladder dysfunction and recent studies of epidural spinal cord stimulation (SCS) accompanied by locomotor training have shown improvements in bladder function^7–9^. In fact, in humans, locomotor training alone improved bladder control^10^, while SCS alone in rats with SCI modulated urethral sphincter activity^11^. Improvements in LUT control may therefore be driven either indirectly or through direct recruitment of afferents innervating the bladder. In the case of limb motion, cervical and lumbar SCS can recruit muscles of the upper and lower limbs^12,13^ by activating reflexes^14,15^, demonstrating that focal stimulation of the afferent system can control motor behaviors^16,17^.

Electrical stimulation of the pelvic and pudendal nerve can produce bladder contractions through a variety of reflex mechanisms^18–22^; stimulating afferents in the pudendal nerve can evoke reflexive micturition^21,23,24^ or suppress ongoing bladder contractions, while afferent activity in the pelvic nerve can modulate this reflex response^25^. These complex reflexes arise in part due to the multiple peripheral targets of the pudendal nerve. The pudendal nerve divides distally into the sensory, deep perineal, and caudal rectal branches^22^. The caudal rectal branch innervates the external anal sphincter and pelvic floor^26^, while the sensory and deep perineal branches innervate the genitalia and urethra. Stimulation of these branches can either reflexively inhibit or evoke micturition^18,22^. However, accessing and instrumenting these nerves could be challenging in humans and will require new surgical procedures, complicating translation of a peripheral nerve-based device^27–30^.

In this study we sought to determine whether epidural SCS can selectively activate the peripheral nerve pathways that control the lower urinary tract. We tested this idea using custom high-density electrode arrays to maximize the opportunity to activate focal sensory inputs while minimizing recruitment of unwanted reflexes. Lower-limb activation frequently accompanies stimulation at the lumbosacral cord^31^, potentially arising from the design of existing SCS arrays that often cover an entire segment of the spinal cord with just a few electrodes^32^. If selective activation were possible, this would establish high-density SCS as a method to directly modulate LUT function by activating sensory afferents in the pudendal and pelvic nerves.

## Results

To test whether high-density SCS could recruit peripheral nerves that innervate the lower urinary tract, we stimulated through each contact on the electrode arrays while they were positioned over the sacral spinal cord and cauda equina (Figure 1). We measured antidromic recruitment of afferents in the pelvic, pudendal and sciatic nerves, and used the relative recruitment of these nerves to determine selectivity in six anesthetized cats. These pathways are co-activated in normal function^18,33^ so we also measured the co-recruitment of all these nerves at increasing stimulation amplitudes.

**Figure 1.**
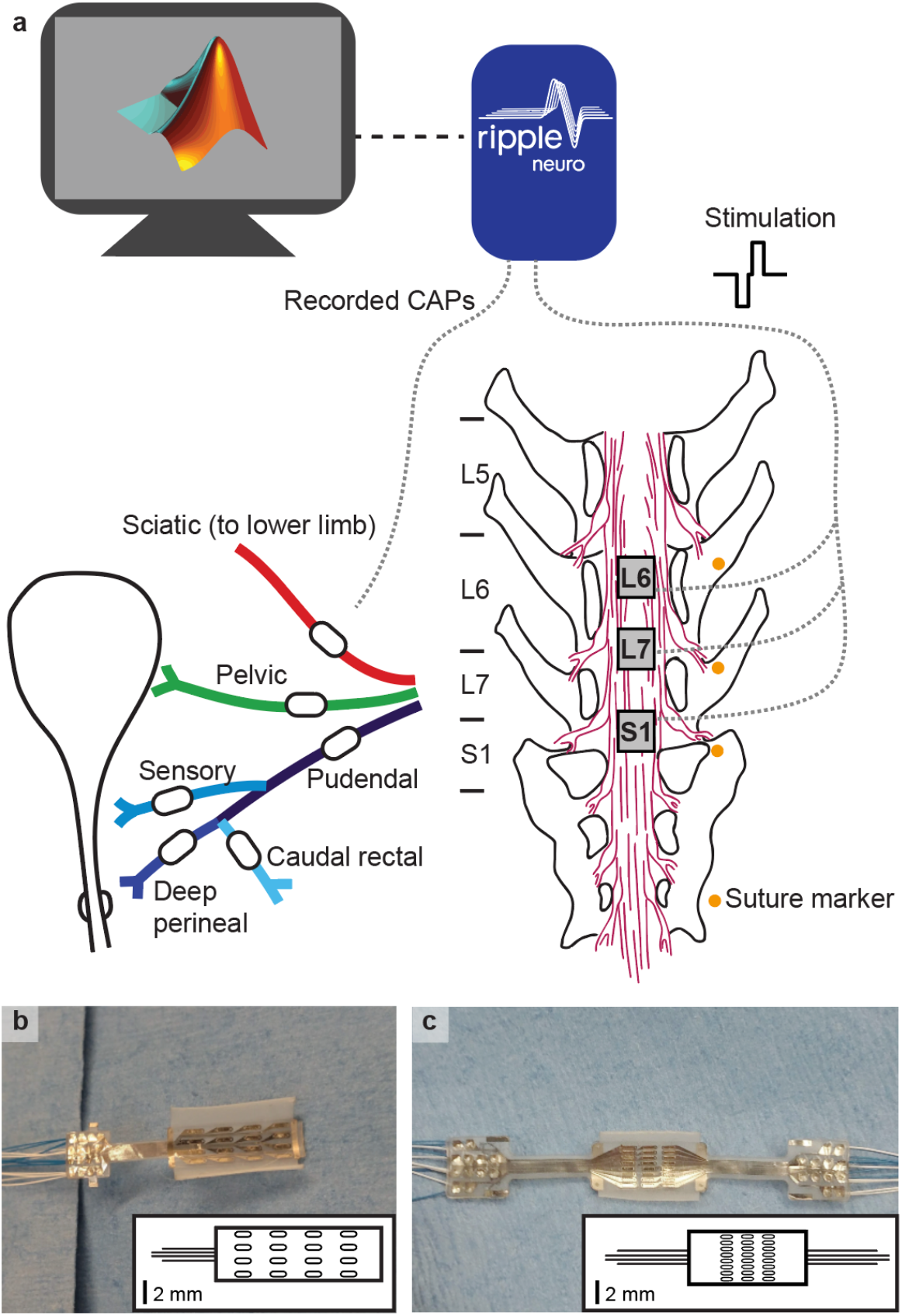
Experimental setup. a) Nerves cuffs, shown as white ovals on each nerve, were placed on multiple peripheral nerves and a high-density electrode array was placed at three locations over the sacral cord and cauda equina. Nerve cuffs on the pelvic nerve (green), pudendal nerve (blue), and pudendal branches (shades of blue) had an inner diameter between 500 μm and 1000 μm. The sciatic nerve (red) cuff had an inner diameter of 3 mm. Recording and stimulation were completed through a MATLAB interface with a Ripple Grapevine system using a closed-loop response detection algorithm. b) In animals 1 and 2, a 16-channel epidural array with four electrode columns spaced laterally across the cord and four electrode rows spaced rostrocaudally was used. The inset shows the array layout to scale, with the wire bundle represented in the same orientation as the photo. The electrodes on the 16-channel array were each 0.45 mm x 1.35 mm and were spaced 0.69 mm apart laterally and 1.64 mm apart rostrocaudally. c) In animals 3-6, a 24-channel epidural array with eight columns and three rows was used. The electrodes on the 24-channel array were each 0.29 mm x 1.0 mm and were spaced 0.23 mm apart laterally and 0.78 apart rostrocaudally.

### High-density SCS selectively recruits pelvic and pudendal afferents

High-density epidural SCS selectively recruited both the pelvic and the pudendal nerves at the sacral cord and cauda equina in all but one animal. Surprisingly, we found that individual electrodes within the array could selectively recruit different nerves even though the electrodes were often less than 1 mm apart. As a typical example (animal 5, Figure 2), an individual electrode selectively recruited the pelvic nerve at 390 μA and the pudendal nerve was not recruited until the amplitude was increased to 460 μA (Figure 2a,b). A nearby electrode recruited the pudendal nerve selectively at 280 μA (Figure 2c).

**Figure 2.**
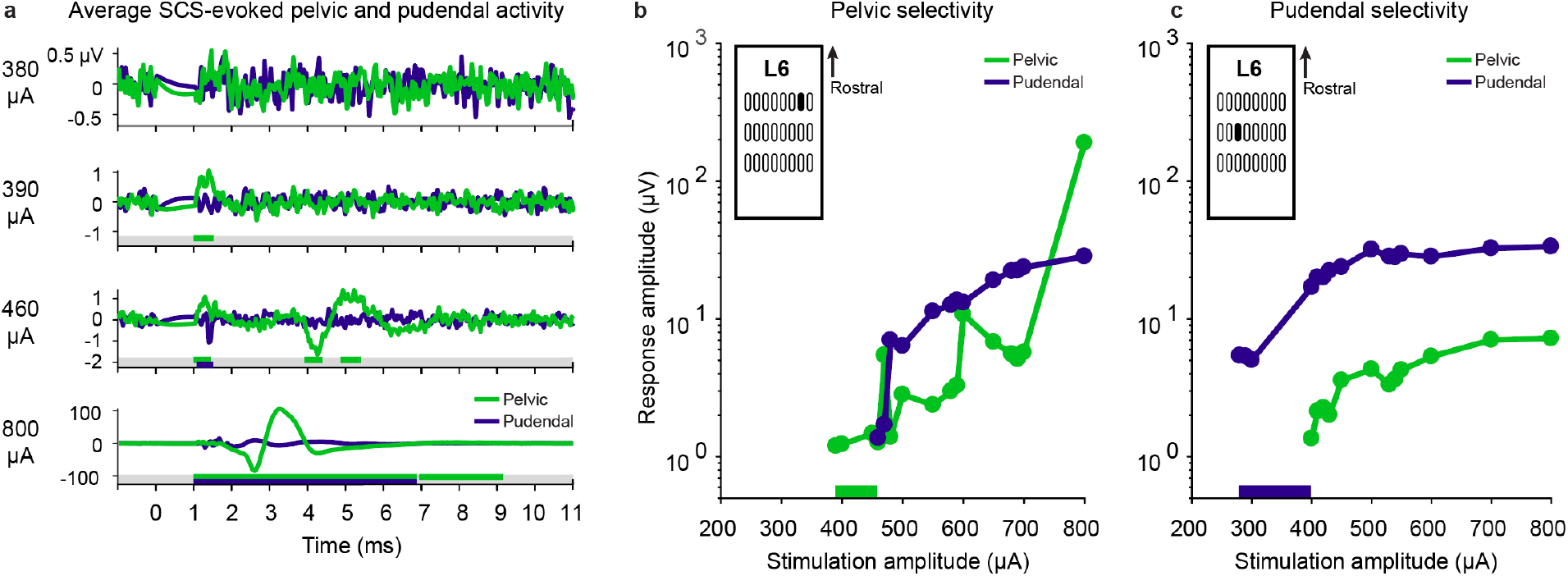
Examples of selective pelvic and pudendal nerve recruitment in animal 5 during stimulation on two different electrodes. a) Stimulation-triggered averages of the pelvic (green) and pudendal (purple) nerve compound nerve action potentials at selected stimulation amplitudes. The traces include 1 ms preceding the stimulus pulse. Windows in which responses were detected are indicated under each trace as colored bars. At 380 μA, no response was detected in either nerve. At 390 μA, a selective response was detected in the pelvic nerve. At 460 μA the pudendal nerve was also recruited. Additional responses in the pelvic nerve at longer latencies also occurred. 800 μA was the maximum stimulation amplitude for this trial and evoked large compound action potentials in the pelvic nerve. Note the different y-axis scales for each stimulation amplitude. b) Peak-to-peak compound action potential amplitude of the pelvic and pudendal nerves for the electrode illustrated in a) that was selective for the pelvic nerve from 390 μA up to 460 μA (selective range highlighted in bar along the x-axis). Only the pudendal and pelvic nerve traces are shown here, but this electrode did not recruit any other instrumented nerves at threshold. The y-axis is shown on a log scale. The specific stimulation electrode is highlighted in the inset. c) Peak-to-peak compound action potential amplitude of the pelvic and pudendal nerves for a nearby electrode (see inset) that selectively recruited the pudendal nerve from 280 μA up to 400 μA (selective range highlighted in bar along the x-axis). The y-axis is shown on a log scale.

We characterized nerve recruitment in three ways: selective recruitment at the threshold amplitude, total recruitment at the threshold amplitude, and recruitment at the maximum stimulation amplitude.

There was no difference in the threshold amplitudes required to selectively recruit the pelvic and pudendal nerves (n=95 selective trials, *p=0.31*, Wilcoxon test), with pelvic nerve thresholds ranging between 150-600 μA and pudendal nerve thresholds ranging between 150-690 μA (Figure 3a, filled circles). Similarly, there was no difference in the threshold amplitude when the pelvic and pudendal nerves were not recruited selectively (n=221 non-selective trials, *p=0.28*, Wilcoxon test) and ranged between 100-800 μA (Figure 3a,b). While there was no difference in the recruitment thresholds between the pelvic and pudendal nerves, when all nerves were considered, there was a significant difference in the threshold amplitudes across subjects (n=6 subjects, *p<0.001*, Kruskal-Wallis test, Figure 3c) and spinal locations (*p<0.001*, Kruskal-Wallis test, Figure 3d). With the arrays placed at the level of the L6 and S1 vertebrae, the threshold amplitudes were lower than with the arrays placed at the L7 vertebra (*p<0.001*, Dunn’s test, Figure 3d) and placing the arrays at the L6 vertebra resulted in lower threshold amplitudes than when they were placed under the S1 vertebra (*p=0.001*, Dunn’s test).

**Figure 3.**
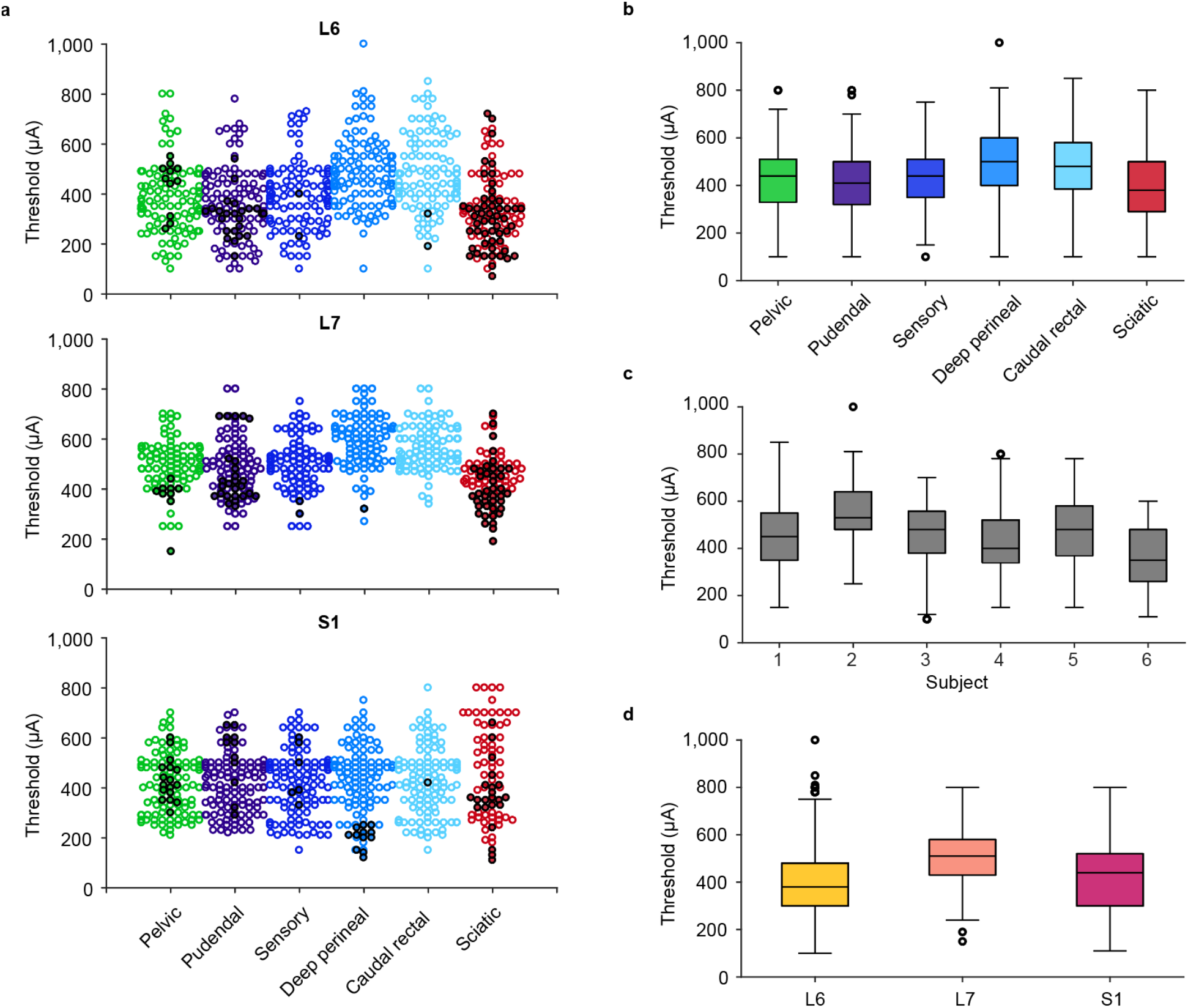
Recruitment thresholds for all animals, nerves and locations. a) Recruitment thresholds for each nerve at each location across all animals. Trials that recruited nerves non-selectively at the threshold amplitude are marked with unfilled circles and trials that recruited nerves selectively at the threshold amplitude are marked with filled circles. b) Recruitment thresholds for each nerve across all locations and animals. c) Recruitment thresholds for each animal across all nerves and locations. d) Recruitment thresholds for each location across all nerves and animals.

The pelvic nerve was recruited selectively at 11 of the 14 tested array locations across the five animals in which selective recruitment occurred. Selective recruitment of the pelvic nerve occurred most often when the array was at the level of the S1 dorsal process (five animals) and occurred on 8.3% of the electrodes (n=368 trials) at this level (Figure 4, dark green bars, Table 1). The pelvic nerve was also recruited selectively in three animals at the L6 and L7 laminar levels (Table 1).

**Fig 4.**
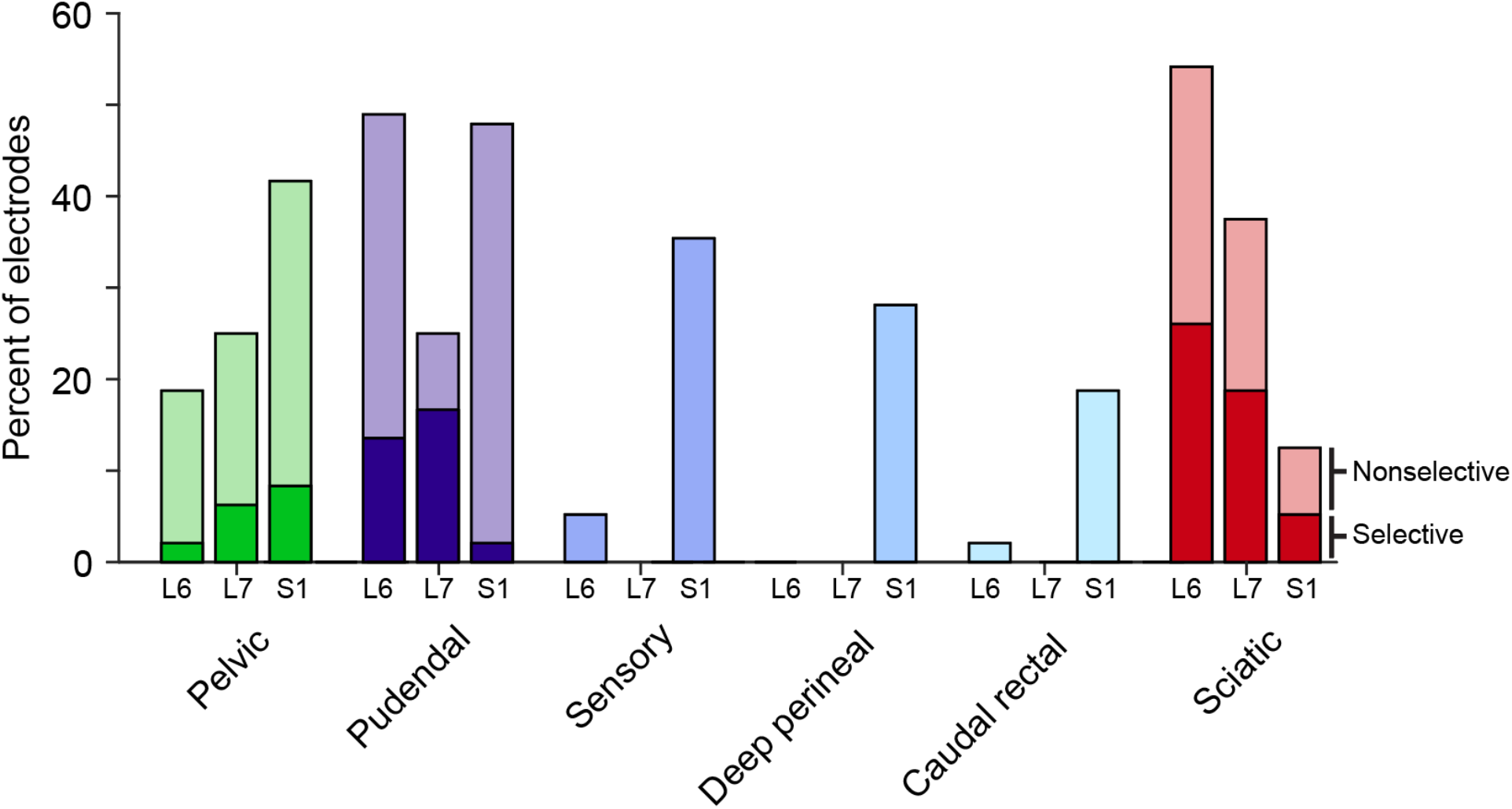
Recruitment of all nerves at threshold at each spinal level. Selective and nonselective nerve recruitment at each location at the threshold amplitude. The darkest color (bottom) in each stacked bar is the median percentage of electrodes that recruited each nerve selectively at the threshold amplitude. The lighter shade (top) represents the percentage of electrodes that recruited each nerve non-selectively at the threshold amplitude. Thus, the cumulative total of the bars represents the total recruitment of each nerve at the threshold amplitude.

**Table 1.** Selective recruitment of each nerve in all animals by vertebral level. The percentage of electrodes at a given array location that selectively recruited each nerve are shown as the median and upper and lower quartiles. The number of animals where stimulation at each array location selectively recruited each nerve is also shown.

| Nerve | L6 Array Placement |  | L7 Array Placement |  | S1 Array Placement |  |
| --- | --- | --- | --- | --- | --- | --- |
|  | % of electrodes | # of animals | % of electrodes | # of animals | % of electrodes | # of animals |
| Pelvic | 2.1 (0.0-16.67) | 3/6 | 6.3 (0.0-16.7) | 3/5 | 8.3 (4.2-18.8) | 5/6 |
| Pudendal | 13.5 (4.2-25.0) | 5/6 | 16.7 (0.0-39.6) | 3/5 | 2.1 (0.0-16.7) | 3/6 |
| Sensory | 0.0 (0.0-4.2) | 2/6 | 0.0 (0.0-3.1) | 1/5 | 0.0 (0.0-12.5) | 2/6 |
| Deep Perineal | 0.0 (0.0-0.0) | 0/6 | 0.0 (0.0-1.0) | 1/5 | 0.0 (0.0-0.0) | 1/6 |
| Caudal Rectal | 0.0 (0.0-0.0) | 1/6 | 0.0 (0.0-0.0) | 0/5 | 0.0 (0.0-0.0) | 1/6 |
| Sciatic | 26.0 (8.3-31.3) | 5/6 | 18.8 (7.3-63.5) | 5/5 | 5.2 (0.0-12.5) | 4/6 |

The pudendal nerve was recruited selectively at 11 of the 14 tested array locations across the five animals in which selective recruitment occurred. Selective recruitment occurred most often with the array at the L6 vertebra (five animals) and occurred on 13.5% of the electrodes at this level (Figure 4, dark purple bars). In 3 animals the pudendal nerve was also recruited selectivity at the lower two levels (Table 1).

On some SCS electrodes the threshold stimulation amplitude evoked activity in multiple peripheral nerves simultaneously. Therefore, we also measured the combined selective and non-selective recruitment at the threshold amplitude. Lastly, we measured nerve recruitment through SCS electrodes at high amplitudes (well above the threshold amplitude) to characterize the maximum recruitment potential of an individual electrode. The pelvic nerve was recruited at threshold on 27.5% of the electrodes across all placements (Figure 4, light green bars, Table 2) and at the maximum stimulation amplitude on 82.3% of the electrodes (see Table 3 for additional detail). The pudendal nerve was recruited at threshold on 41.3% of the electrodes across all placements (Figure 4, light purple bars, Table 2) and at the maximum amplitude on 84.2% of electrodes (see Table 3 for additional detail). On 14.0% of the electrodes, or about 3-4 electrodes on a 24-channel array, stimulation at maximal intensities evoked no detectable response on either the pelvic or pudendal nerves.

**Table 2.** Total threshold recruitment of each nerve in all animals by vertebral level. The percentage of electrodes at a given array location that recruited each nerve selectively or non-selectively at threshold are shown as the median and upper and lower quartiles. The number of animals where stimulation at each array location recruited each nerve is also shown.

| Nerve | L6 Array Placement |  | L7 Array Placement |  | S1 Array Placement |  |
| --- | --- | --- | --- | --- | --- | --- |
|  | % of electrodes | # of animals | % of electrodes | # of animals | % of electrodes | # of animals |
| Pelvic | 18.8 (12.5-29.2) | 5/6 | 25.0 (6.3-34.4) | 4/5 | 41.7 (29.2-75.0) | 6/6 |
| Pudendal | 49.0 (33.3-56.3) | 5/6 | 25.0 (14.1-63.5) | 4/5 | 47.7 (41.7-68.8) | 5/6 |
| Sensory | 5.2 (0.0-8.3) | 4/6 | 0.0 (0.0-21.9) | 2/5 | 35.4 (20.8-68.8) | 6/6 |
| Deep Perineal | 0.0 (0.0-4.2) | 2/6 | 0.0 (0.0-1.0) | 1/5 | 28.1 (12.5-62.5) | 6/6 |
| Caudal Rectal | 2.1 (0.0-4.2) | 3/6 | 0.0 (0.0-1.0) | 1/5 | 18.8 (8.3-31.3) | 6/6 |
| Sciatic | 54.2 (41.7-75.0) | 6/6 | 37.5 (19.8-63.5) | 5/5 | 12.5 (0.0-12.5) | 4/6 |

**Table 3.** Maximum amplitude recruitment of each nerve in all animals by vertebral level. The percentage of electrodes at a given array location that recruited each nerve at the maximum amplitude are shown as the median and upper and lower quartiles. The number of animals where stimulation at each array location recruited each nerve is also shown.

| Nerve | L6 Array Placement |  | L7 Array Placement |  | S1 Array Placement |  |
| --- | --- | --- | --- | --- | --- | --- |
|  | % of electrodes | # of animals | % of electrodes | # of animals | % of electrodes | # of animals |
| Pelvic | 83.3 (75.0-87.5) | 6/6 | 91.7 (80.2-94.3) | 5/5 | 83.3 (62.5-100.0) | 6/6 |
| Pudendal | 87.5 (62.5-91.7) | 6/6 | 91.7 (84.4-96.9) | 5/5 | 81.3 (66.7-100.0) | 6/6 |
| Sensory | 87.5 (62.5-91.7) | 6/6 | 91.7 (78.1-96.9) | 5/5 | 83.3 (66.7-100.0) | 6/6 |
| Deep Perineal | 87.5 (50.0-95.8) | 6/6 | 87.5 (72.9-89.6) | 5/5 | 81.3 (58.3-100.0) | 6/6 |
| Caudal Rectal | 87.5 (33.3-91.7) | 6/6 | 83.3 (75.5-92.7) | 5/5 | 81.3 (58.3-100.0) | 6/6 |
| Sciatic | 83.3 (75.0-87.5) | 6/6 | 91.7 (80.2-94.3) | 5/5 | 83.3 (62.5-100.0) | 6/6 |

### Recruitment of pudendal nerve branches

Activating different branches of the pudendal nerve can have different effects on bladder function^22,24^, and we were therefore interested in determining the recruitment properties of these individual branches (sensory, deep perineal, and caudal rectal). SCS evoked responses in every branch of the pudendal nerve in every cat at the maximal stimulation amplitude (Table 3). There was no difference in the number of electrodes that could recruit these nerves at different spinal levels at these high intensities (*p=0.989*, Kruskal-Wallis test).

Pudendal nerve branches were recruited at the threshold amplitude most often at the S1 location (Figure 4, blue bars). At this location, all three branches were recruited at threshold in every animal (Table 2) and the caudal rectal and deep perineal nerves were significantly more likely to be recruited at threshold compared to other locations (*p<0.005,* Kruskal-Wallis test), although this difference was not significant for the sensory branch (*p=0.069*, Kruskal-Wallis test).

Selective recruitment of the pudendal nerve branches was rare in all animals at all spinal locations (Table 1). The sensory branch was recruited selectively at just five of the 17 total tested locations. The other two branches were each selectively recruited at only two of the 17 locations, with the deep perineal branch recruited selectively on just 13 electrode contacts across all experiments and the caudal rectal branch recruited selectively on just three contacts.

Lastly, the stimulation amplitude required to recruit the deep perineal and caudal rectal nerves was significantly higher than all other nerves at the L6 and L7 laminar levels (n=97 L6 trials and 93 L7 trials, *p<0.001*, Kruskal-Wallis test, Figure 3a. This was not true at the S1 lamina level (n=97 trials, *p=0.360*, Kruskal-Wallis test, Figure 3a).

### Sciatic nerve recruitment

A common side effect of electrically stimulating the sacral cord or nerves is lower limb movement resulting from sciatic nerve activation^7,11,34,35^. Therefore, we monitored sciatic nerve activity during these experiments and found that similar to the results for the pelvic and pudendal nerves, SCS activated the sciatic nerve both selectively and non-selectively at all levels. The sciatic nerve had lower thresholds than all nerves except for the pudendal nerve at the L6 location (n=113 trials, *p<0.001,* Kruskal-Wallis test) and lower thresholds than all nerves at the L7 location (n=101 trials, *p<0.001,* Kruskal-Wallis test). However, sciatic threshold amplitudes were no different than the other nerves at the S1 lamina location (n=106 trials, *p=0.360*, Kruskal-Wallis test).

These lower thresholds compared to LUT nerves are likely responsible for the fact that at the L6 and L7 laminar locations, the sciatic nerve was recruited selectively more often than any LUT nerve (Figure 4, dark red bars). However, at the S1 location, there was no difference in the number of electrodes that selectively recruited LUT and sciatic nerves (*p=0.185*, Kruskal-Wallis test).

### Nerve selectivity changes with stimulation location

In most animals the pelvic, pudendal, and sciatic nerves were selectively recruited by multiple electrodes. We investigated the extent to which these patterns of recruitment tended to be organized within the array. Figure 5a shows a representative example of the nerves selectively recruited by individual electrodes across the three array levels in one animal. Many electrodes recruited nerves selectively (Figure 5a, colored rectangles), while other electrodes only recruited nerves non-selectively (Figure 5a, black rectangles). No obvious organization was seen when we considered only purely selective electrodes. However, when we examined both selective and non-selective responses at threshold, we observed stimulation ‘hot spots’ within the arrays (Figure 5b). To quantify similarities in spatial recruitment we compared the nerves recruited on neighboring electrodes to the nerves recruited by each individual electrode. When we stimulated through an electrode that activated a particular nerve at threshold, 62.5% (IQR 38.2-80.0%) of the neighboring electrodes recruited that same nerve at threshold (n=320 trials). Conversely, if an electrode did not recruit a nerve at threshold, the neighboring electrodes were also unlikely to recruit that nerve at threshold, with a median recruitment of 0.0% (IQR 0.0-20.0%) (n=318 trials, Figure 5c, gray bars). Even though individual electrodes were very close to each other (0.23-0.78 mm), 37.5% (IQR 0.0%-62.5%) of adjacent electrodes recruited at least one different nerve. Every nerve was recruited more frequently at threshold when it was also recruited at threshold on an adjacent electrode (Figure 5c, colored bars).

**Figure 5.**
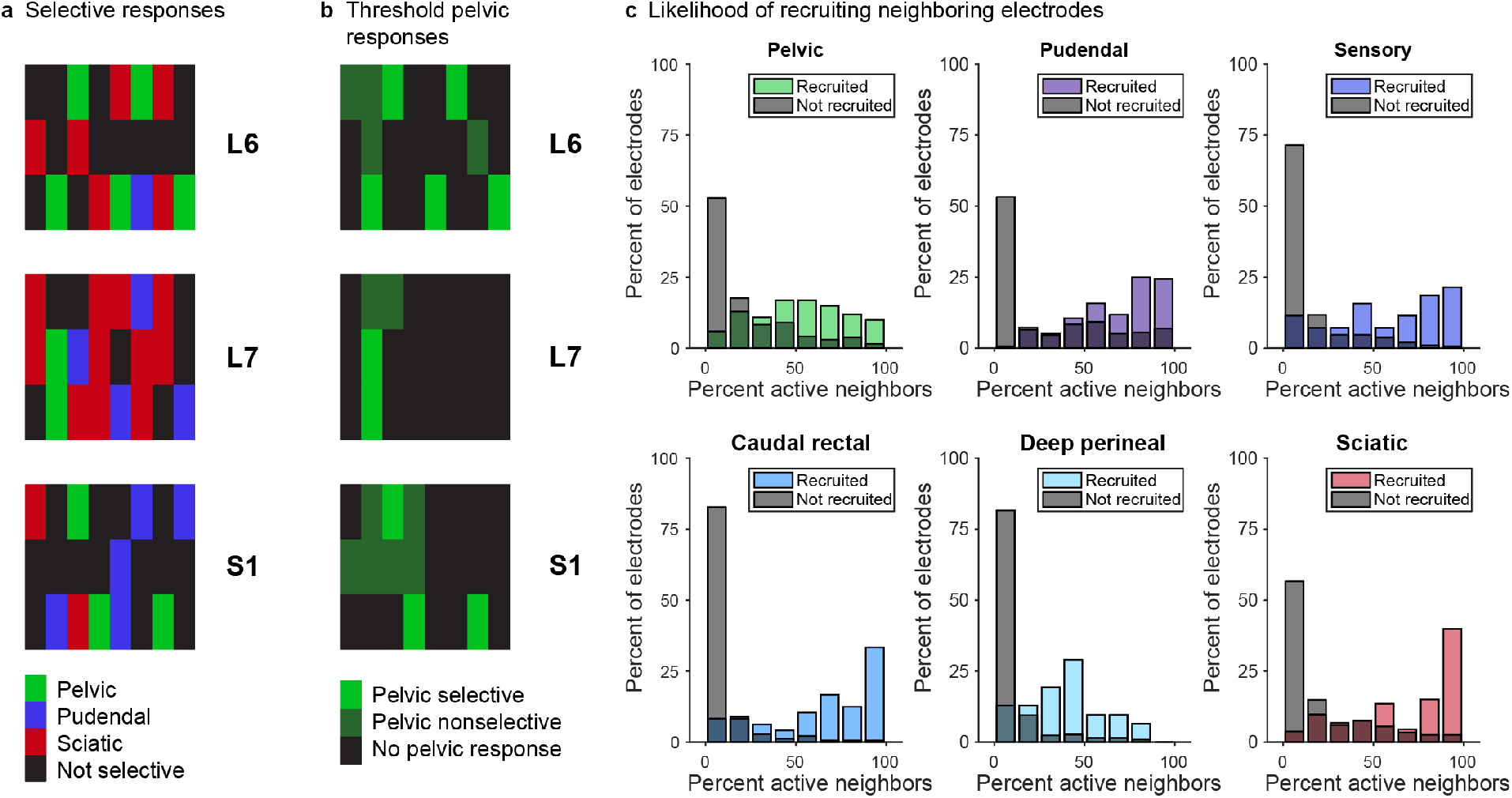
Spatial arrangement of evoked responses. a) Selective recruitment for the pelvic nerve, pudendal nerve, and sciatic nerve in a representative animal. b) Pelvic nerve recruitment for the same animal as panel a, demonstrating that the pelvic nerve was frequently recruited non-selectively on electrodes adjacent to selective electrodes. c) When an electrode activated a particular nerve at the threshold amplitude, neighboring electrodes were likely to activate that nerve as well (colored bars). The y-axis is normalized to the total number of electrodes that activated a specific nerve. When that nerve had not been activated, surrounding electrodes were much less likely to activate neighboring electrodes (gray bars). The y-axis for the gray bars is normalized to the total number of electrodes that did not activate a specific nerve.

This relationship occurred not only at threshold, but was also true at the maximum stimulation amplitude. At the maximum amplitude, if an electrode recruited a nerve, 84% of surrounding electrodes also recruited that nerve (n=320 trials). However, if an electrode did not recruit a nerve, only 51% of surrounding electrodes did (n=149 trials).

### Nerve coactivation

Because we observed numerous electrode contacts with non-selective nerve recruitment, we wanted to quantify the extent to which this recruitment was limited to LUT nerves as compared to co-activation with the off-target sciatic nerve. This co-activation of different groups of nerves varied by level.

At the L6 and L7 levels, the sciatic nerve was coactivated with the LUT nerves on most occasions (Fig. 6a,b). Given the overall lower recruitment of the sciatic nerve at the S1 lamina level, coactivation was much less common at this level (Fig. 6c). Conversely, the pelvic and pudendal nerves were coactivated more frequently at the S1 level (Fig. 6c) than at the two more rostral spinal locations (Figure 6a,b). When the pudendal nerve was recruited, the pelvic nerve was also active 30.3%, 33.3% and 69.3% of the time at the L6, L7, and S1 locations respectively.

**Figure 6.**
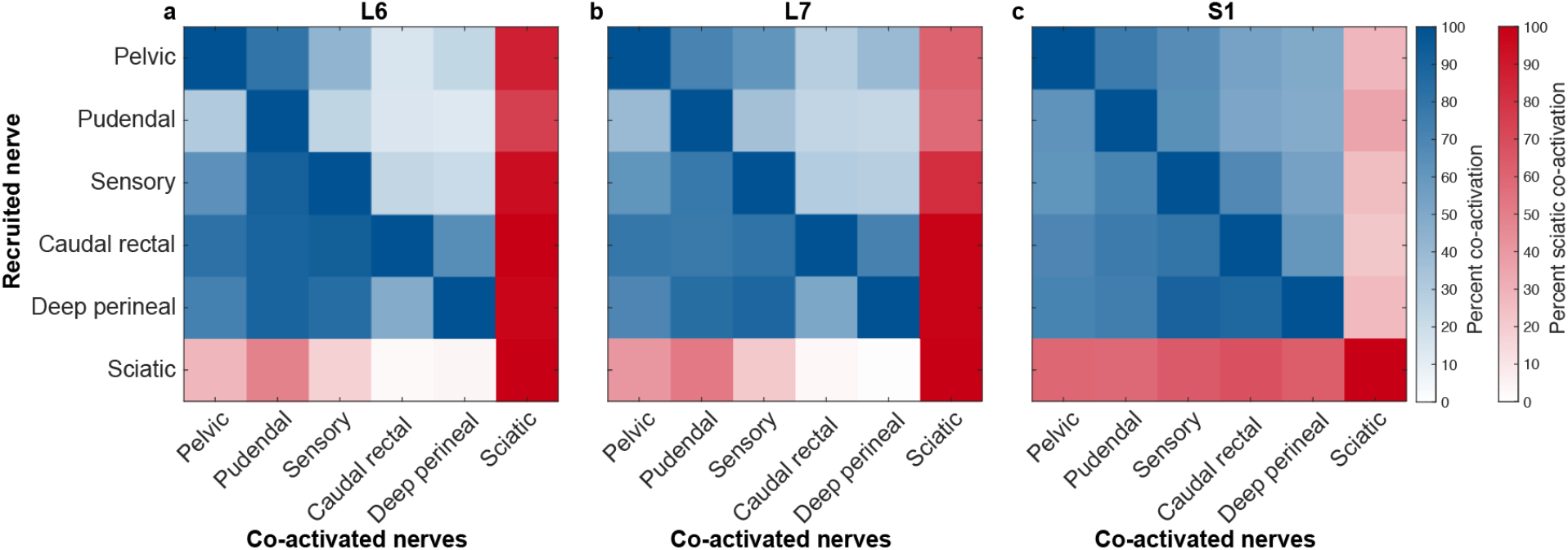
Coactivation of all nerves. When a nerve first became active (vertical axis), other nerves are often co-activated or were recruited at lower amplitudes (horizontal axis). Sciatic comparisons are colored differently for clarity.

### Dynamic range of recruited nerves

While the primary aim of this experiment was to identify whether peripheral afferents could be recruited selectively by SCS, we also wanted to characterize the dynamic range of stimulation on each electrode. The dynamic range is the stimulus amplitude range between the threshold amplitude and the stimulation amplitude at which additional nerves are recruited. Within this range, stimulation remains selective. The dynamic range was typically smallest with the array placed at the S1 lamina level (*p<0.001*, Kruskal-Wallis test, Figure 7) and had a median of 20 μA across all electrodes and animals. With the array placed at the L6 and L7 laminar levels, the median dynamic range was 50 μA.

**Figure 7.**
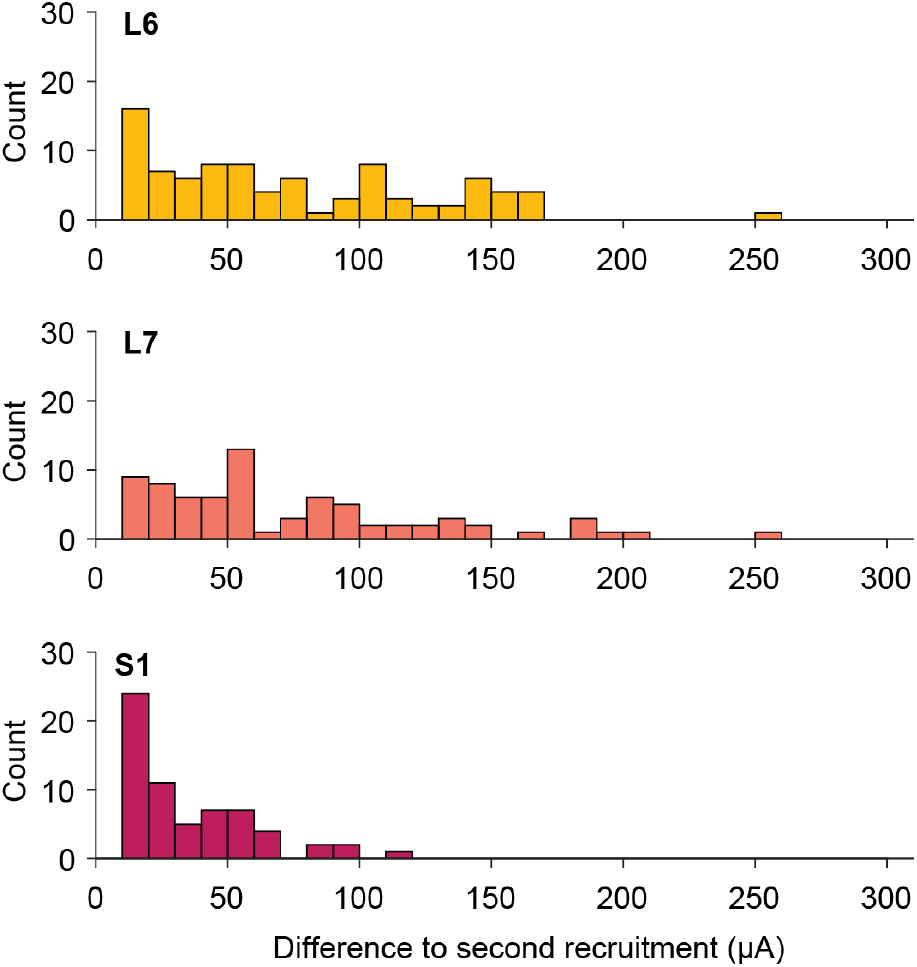
Distribution of dynamic ranges for each array location. The dynamic range of each selective electrode is the amount of additional stimulation current necessary to evoke activity in an additional nerve, over and above the initial selective response. Many nerves recruited selectively had a dynamic range of less than 50 μA.

The dynamic range also varied between different nerve groups. The amplitude difference between LUT nerve recruitment and coactivation with other LUT nerves was 20 μA (IQR 10-30 μA) (n=149 trials). For LUT nerves to become coactivated with the sciatic nerve, the dynamic range was slightly higher at 30 μA (IQR 10-50 μA) (n=47 trials). Finally, when the sciatic nerve was recruited selectively, an amplitude increase of 80 μA (IQR 40-140 μA) was required to recruit LUT nerves (n=115 trials).

## Discussion

We found that epidural SCS over the sacral cord can recruit afferent axons arising from the pelvic and pudendal nerves, providing a mechanism to directly modulate bladder function. While completely selective recruitment of LUT nerves was not common (Fig. 4), selective recruitment of the pelvic, pudendal, and sciatic nerves was possible at all three spinal levels tested and in five of the six animals.

Epidural SCS has been explored in combination with locomotor training to improve lower urinary tract control in humans^7^ and rats^8^. However, it was unclear from these studies whether the benefits of stimulation on bladder function arose directly from SCS itself, or whether the benefits were primarily driven by indirect effects such as improved mobility. Our results demonstrate a plausible physiological mechanism for SCS to directly recruit LUT reflexes and modulate bladder function. In these experiments, both pelvic and pudendal nerves were activated by SCS and stimulating pelvic nerve^36,37^ and pudendal nerve ^18,21–23^ afferents can facilitate or inhibit micturition. While it might seem more obvious to directly target the pelvic and pudendal nerves for stimulation, surgical access to these nerves can be challenging in humans, and the pelvic nerve is particularly inaccessible^38,39^. Here, we demonstrate that it is possible to access LUT afferent pathways using sacral SCS, although the degree to which it is important to selectively activate specific branches remains unclear.

We have further demonstrated in this study that high-density spinal cord arrays are able to produce completely different recruitment patterns within the same spinal level simply by changing the active electrode within the array. This result suggests that high-density electrode arrays could be particularly beneficial to optimize SCS for improving bladder function–or any other function of interest at other spinal levels^40–45^–and that existing commercial stimulation leads may be inadequate.

We placed our stimulation electrodes over the sacral spinal cord rather than more rostral levels of the cord, where many clinical implants are placed^46–49^ as the sacral cord contains the motoneuron pools of the bladder and urethral sphincter^50^ and gives rise to the entirety of the pelvic and pudendal nerves^51–53^. While there were some differences between stimulation locations in terms of threshold amplitudes (Figure 3d), selectivity (Figure 4) and dynamic range (Figure 7), these differences were subtle. In fact, within any given animal, we were able to recruit all instrumented nerves at all levels.

We evoked the most activity in pelvic afferents when the array was at more caudal locations while pudendal nerve afferents were recruited at more rostral locations (Figure 4). This is consistent with previous anatomical observations of the roots that contribute to each of these nerves. The pudendal nerve is typically composed of fibers from the S1 and S2 roots, while the pelvic nerve is typically composed of fibers from the S2 and S3 roots^54,55^. Afferents of the pelvic nerve may however be located more rostrally, in the S1 and S2 roots, in some animals^56^. Furthermore, motoneuron pools for muscles innervated by the pudendal nerve tend to be located in the S1 and S2 cord in Onuf’s nucleus, while motoneuron pools for muscles innervated by the pelvic nerve are typically focused in the S2 and S3 cord^50,55^.

### Lower limb activation

If the sciatic nerve were always activated during stimulus trains intended to recruit LUT nerves, the associated lower-limb movement could be very disruptive. In fact, activation of the lower limb is a common problem with the commercially available InterStim sacral nerve stimulators^57^ and the Finetech anterior root stimulation system^58^. Although motor activation of the lower-limb does not prevent bladder prostheses from being effective, it is typically an undesirable off-target effect^7,11,34,35^. On the other hand, recruiting sensory afferents of the lower limb, particularly the tibial nerve, has been shown to improve continence^59^, making this a potentially useful target in some contexts. We found that in the spinally intact cat, there was substantially less activation of the sciatic nerve when the electrodes were over the cauda equina compared to the sacral cord, which is consistent with the path of the lower lumbar and sacral roots within the spinal canal at these locations.

### LUT co-activation

While this study focused on selective nerve activation, axons in many different LUT nerve branches are active simultaneously in behavior. For instance, the anal sphincter, innervated by the caudal rectal nerve, and the external urethral sphincter, innervated by the deep perineal nerve, are frequently coupled^33^. Further, in some cases, co-stimulation of multiple pudendal branches improves voiding efficiency^24^. It is therefore likely that selectively activating these branches may not be necessary to effectively control bladder function. This is encouraging because we found that the dynamic range for selective stimulation was typically less than 100 μA.

The sensory branch typically had lower threshold amplitudes than the deep perineal and caudal rectal branches (Figure 4b). Because SCS primarily recruits afferents, this difference in threshold could be due to the high density of afferent fibers in the sensory branch compared to both the deep perineal and caudal rectal branches, which have substantial motor functions^54^.

### Limitations

In this study we focused on selective recruitment of individual nerves. While we saw that it was possible to selectively activate most nerves in most animals, the actual number of electrode contacts that selectively recruited LUT nerves was low. However, this selectivity may not be necessary, as many functional behaviors require coactivation of multiple pathways. If improved selectivity were directly related to functional control of the LUT, it would be useful to maximize the number of electrode contacts that selectively recruited LUT afferents.

In this study we used monopolar stimulation exclusively, which allowed us to systematically test all the electrodes in the available time, but may have been suboptimal to recruit afferent populations selectively. To improve selectively, multipolar stimulation can be used to localize or focus current between several electrodes^60^. In a recent study in humans, multipolar stimulation was often required to evoke meaningful sensory percepts in amputees, while monopolar stimulation was generally less effective^61^. Similarly, some commercially available SCS systems leverage multipolar stimulation to increase the focality of the paresthesias evoked by stimulation^62^. Additional selectivity could potentially be gained by changing stimulation waveforms^63,64^ or applying variable-frequency stimuli^65^.

Another limitation of this study is that it was conducted in acute experiments and we did not determine the stability of these effects during movement. Postural effects are known to be considerable in human spinal cord stimulation^47^, and these effects could be exacerbated using small electrodes of the type considered here. Additionally, this study was performed in cats, and the reduced cerebrospinal fluid thickness compared to humans may impact parameter choice, particularly threshold amplitudes. The smaller spinal cord size in cats also requires fewer electrodes to cover the spinal cord area. The large number of electrodes that could be required in a human application might also require new methods of parameter tuning to be developed, including closed-loop methods where muscle activity or other non-invasively accessible signals are used to automatically tune parameters. Ultimately, the goal of this work is to manipulate LUT function, and here we only study peripheral nerve recruitment, particularly recruitment of the sensory fibers. Future studies will expand this work to include direct measures of LUT function in response to stimulation.

### Implications for neuroprosthetic devices

This study demonstrates that it is possible to selectively activate individual peripheral nerves innervating the LUT with high-density SCS. This understanding could potentially provide a route to improve upon recent studies where results may vary considerably between individuals^7^, as it illuminates the variability in recruitment that could occur with subtly changing electrode positions. This study therefore supports the design and development of new high-density electrodes to achieve selective activation, which may improve the effects of human SCS trials. Finally, this study highlights the potential use of epidural SCS to target autonomic systems generally^66^ by adding a physiological and scientific basis for stimulating these pathways.

## Methods

### Surgical procedures

Acute experiments were conducted under isoflurane anesthesia in 6 adult male cats weighing between 4.1 and 6.4 kg. All procedures were approved by the University of Pittsburgh Institutional Animal Care and Use Committee. All procedures were performed in accordance with the relevant guidelines and regulations.

The animals were anesthetized with a ketamine/acepromazine cocktail and anesthesia was maintained using isoflurane (1-2%). A tracheostomy was performed and the trachea was cannulated and connected to an artificial respiration system. Throughout the procedure, the animal was artificially ventilated at 12-14 breaths per minute. Blood pressure was monitored with a catheter placed in the carotid artery. Temperature was maintained with a warm air heating pad and IV fluids were administered continuously. End tidal CO_2_, SpO2, core temperature, heart rate and blood pressure were monitored throughout the procedure and kept within a normal physiological range. Following experimental data collection, animals were euthanized with an IV injection of Euthasol.

The bladder was exposed through a midline abdominal incision and a dual-lumen catheter (Model CDLC-6D, Laborie Medical Technologies, Williston, VT) was placed through the bladder dome to measure bladder pressure as well as to infuse and withdraw fluids. The catheter was secured in place with a purse string suture.

To measure antidromic compound action potentials evoked by spinal cord stimulation, we placed bipolar nerve cuffs (Micro-Leads Inc., Somerville, MA) on the left pelvic and pudendal nerves as well as pudendal nerve branches (Figure 1a). The pelvic nerve was dissected free near the internal iliac artery and a cuff was placed prior to the branching of the pelvic plexus. The abdominal incision was then closed in layers and the animal was placed in the prone position. We made an incision on the left hindquarters between the base of the tail and the ischial tuberosity and performed blunt dissection to expose the pudendal nerve. We then placed nerve cuffs on the left pudendal nerve and the sensory, deep perineal, and caudal rectal branches^22^ of the left pudendal nerve. We also placed a five-pole spiral nerve cuff (Ardiem Medical, Indiana, PA) on the left sciatic nerve to measure off-target effects associated with the lower-limb. A recording reference electrode, consisting of a stainless steel wire with ~1 cm of insulation removed, was placed subcutaneously in the left lower back.

We performed a laminectomy at the L6, L7, and S1 vertebral levels to expose the sacral spinal cord, then placed a custom epidural spinal cord array (Micro-Leads Inc., Somerville, MA) with 16 or 24 channels (Figure 1b,c) on the spinal cord at three different locations over the sacral spinal cord and cauda equina (Figure 1a). The electrodes on the 16-channel array were each 0.45 mm x 1.35 mm and were spaced 0.69 mm apart laterally and 1.64 mm apart rostrocaudally (Figure 1b, inset shown to scale). The electrodes on the 24-channel array were each 0.29 mm x 1.0 mm and were spaced 0.23 mm apart laterally and 0.78 apart rostrocaudally (Figure 1c, inset shown to scale). The centers of the L6, L7 and S1 dorsal spinal processes were marked using a suture placed in paraspinal muscles prior to the laminectomy. The epidural arrays were then placed on the epidural surface of the spinal cord and aligned to these suture markers. For the most rostral location, the arrays were placed such that the most rostral electrode on the array was aligned with the center of the L6 dorsal process. After experiments were completed at this location the electrode array was moved so that the most rostral electrode on the array was aligned with the L7 suture marker. For the final location, the most caudal electrode on the array was aligned with the S1 suture marker. A stimulation return electrode, consisting of a stainless steel wire with ~1 cm of insulation removed, was placed outside the spinal column, near the L7 transverse process. Landmarks and dorsal root entry zones were verified postmortem.

Five animals were tested at three spinal levels and one animal tested at two spinal levels (L6 and S1), giving a total of 17 sets of data.

### Neural recording and stimulation

Stimulation was delivered with a Grapevine Neural Interface Processor through a Nano 2+ Stim high-current headstage (Ripple LLC), with stimulation patterns commanded from MATLAB (MathWorks Inc, Natick, MA). This headstage delivers stimulation current amplitudes of up to 1.5 mA and has a compliance voltage of ±8.5 V. The stimulation amplitude across all trials ranged from 10-1500 μA with a resolution of 10 μA between steps up to 1280 μA, and a resolution of 20 μA from 1280-1500 μA. Stimulation pulses were symmetric with 200 μs cathodal and anodal phases. Phases were separated by a 66 μs interphase interval. For animals 1-2 and 5-6, the cathodal phase was applied first, followed by the anodal phase. For animals 3-4, the anodal phase was applied first. Regardless of which phase was applied first, the recruitment thresholds were no different (*p=0.28,* Wilcoxon test).

Compound action potentials were sampled at 30 kHz with a Surf S2 headstage (Ripple LLC) through the Grapevine Neural Interface Processor. The signal was filtered with a high-pass filter with a 0.1 Hz cutoff followed by a low-pass filter with a 7.5 kHz cutoff, using 3^rd^ order Butterworth filters. The signals from each pole of the bipolar nerve cuffs were then differenced to find the response on a given nerve.

### Compound action potential detection

Stimulation artifacts were removed from the nerve cuff recordings by linearly interpolating between the sample immediately before the onset of each stimulus pulse to 0.5 ms after the end of each stimulus pulse. We then high-pass filtered the signal at 300 Hz using a 2^nd^ order Butterworth filter. The signal-to-noise ratio for detecting antidromic action potentials at the recruitment threshold is substantially less than one due to the presence of spontaneous activity in the nerves as well as general recording noise. Therefore, we used stimulus-triggered averaging to detect responses evoked by stimulation. The presence of a compound action potential on each nerve was determined by comparing responses following stimulation to baseline recordings in which no stimulation occurred, using a previously-published method^67,68^. To determine the response detection threshold, we calculated the 99% confidence interval on the root mean squared baseline amplitude. We then set the detection threshold to 3.2 standard deviations above the upper bound of this baseline mean, or a minimum of 0.5 μV. This threshold was determined empirically to most accurately detect true responses without false positives. The stimulus-triggered average was calculated 200 times from a random subsample of 80% of the responses in order to find a distribution of typical responses. In each of these responses, the root mean squared amplitude of each time window was compared to the root mean squared of the baseline amplitude, using a 250 μs sliding window with a 25 μs overlap. If 95% of these responses were suprathreshold and nerve activity was detected for at least three consecutive windows, the response was considered significant.

### Determining recruitment thresholds

A binary search procedure was used to determine the minimum stimulus current necessary to recruit each nerve according to methods published previously^67,68^. First, we delivered 50 stimulation pulses through each electrode on the array in a random order using high amplitude pulses at 20 Hz to determine which electrodes could evoke responses in the peripheral nerves. The stimulation amplitude for this trial was determined based on the highest amplitude that did not evoke substantial movement in the leg, or when all electrodes showed neural responses, and typically ranged from 600-1000 μA. After the responses to stimulation at the maximum amplitude were determined, all stimulation electrodes that evoked compound action potentials in at least one instrumented nerve were tested individually using a binary search procedure to determine the thresholds for every nerve recruited. We set the stimulation frequency during these trials by determining the longest-latency neural response on each nerve cuff, adding 5 ms, and taking the inverse of this time. With this approach, we were able to maximize the stimulation frequency and minimize overall experiment time. 300 stimulation pulses were delivered to each electrode at each tested amplitude. For each nerve showing a response, we determined the current threshold to a resolution of 10 μA. This procedure typically took 2-3 hours for each location of the spinal cord array.

To determine the selectivity of this high-density SCS electrode, we measured the recruitment thresholds, selectivity, and dynamic range of stimulation-evoked neural responses. Recruitment thresholds for each nerve were defined as the lowest amplitude at which a response to SCS was detected. Threshold responses were considered to be selective when only a single nerve responded at the threshold amplitude and non-selective when multiple nerves were simultaneously recruited at the threshold amplitude. We determined whether pudendal nerve branches were activated selectively by excluding the pudendal nerve and comparing their recruitment thresholds only to other branches, the pelvic nerve, and the sciatic nerve. We defined the dynamic range of stimulation on an electrode as the difference between the threshold amplitude for the recruited nerve and the first higher amplitude at which multiple nerves were recruited.

### Statistics

The recruitment thresholds were not normally distributed (*p < 0.001*, Lilliefors test) so data are reported as the median threshold amplitude with lower and upper quartiles. The Wilcoxon rank-sum test was used to test for differences between two groups. For comparisons between multiple groups of data, we used a Kruskal-Wallis test with a Dunn’s test for post-hoc analysis. The data were analyzed in Matlab 2018a (Mathworks, Natick, MA).

## Data Availability

All data collected for this study and used in these analyses are available at https://doi.org/10.26275/iami-zirb.^69^ This dataset also contains data from other experiments, so the animals described in this study are referred to as subjects 54, 60, 64, 63, 68, 69, and 78 in the available dataset, and this study only includes monopolar stimulation data. These data are provided using the CC BY 4.0 license.

## Acknowledgements

Research reported in this publication was supported by the Office Of The Director, National Institutes Of Health of the National Institutes of Health under Award Number OT2OD024908. The content is solely the responsibility of the authors and does not necessarily represent the official views of the National Institutes of Health. We thank the veterinary staff at Magee-Womens Research Institute for their help with animal care and experiments, as well as Massimiliano Novelli for his help with data management and Ameya Nanivadekar for his advice on data analysis.

## Author Contributions

RAG and BLM conceived of the overall project. MKJ, CHG, LEF and RAG designed the experiments. MKJ, CHG, RK and RAG collected the data. LW, JIO, and CC designed and manufactured the custom electrode arrays. MKJ analyzed the data. MKJ and RAG wrote the manuscript. All authors contributed to the interpretation of the data and provided critical review and approval of the manuscript.

## Competing Interests

BLM, LW, CC, and JIO are employees of Micro-Leads Inc. who design and develop implantable electrodes. The other authors declare no competing interests.

## Notes

https://doi.org/10.26275/iami-zirb

